# AlphaFamImpute: high accuracy imputation in full-sib families from genotype-by-sequencing data

**DOI:** 10.1101/2019.12.11.872432

**Authors:** Andrew Whalen, Gregor Gorjanc, John M Hickey

**Affiliations:** The Roslin Institute and Royal (Dick) School of Veterinary Studies, The University of Edinburgh, Midlothian, Scotland, UK

## Abstract

**Summary:** AlphaFamImpute is an imputation package for calling, phasing, and imputing genome-wide genotypes in outbred full-sib families from single nucleotide polymorphism (SNP) array and genotype-by-sequencing (GBS) data. GBS data is increasingly being used to genotype individuals, especially when SNP arrays do not exist for a population of interest. Low-coverage GBS produces data with a large number of missing or incorrect naïve genotype calls, which can be improved by identifying shared haplotype segments between full-sib individuals. Here we present AlphaFamImpute, an algorithm specifically designed to exploit the genetic structure of full-sib families. It performs imputation using a two-step approach. In the first step it phases and imputes parental genotypes based on the segregation states of their offspring (that is, which pair of parental haplotypes the offspring inherited). In the second step it phases and imputes the offspring genotypes by detecting which haplotype segments the offspring inherited from their parents. With a series of simulations we find that AlphaFamImpute obtains high accuracy genotypes, even when the parents are not genotyped and individuals are sequenced at less than 1x coverage.

**Availability and implementation:** AlphaFamImpute is available as a Python package from the AlphaGenes website, http://www.AlphaGenes.roslin.ed.ac.uk/AlphaFamImpute.

**Supplementary information:** A complete description of the methods is available in the supplementary information.

## Introduction

AlphaFamImpute is a software package for calling, phasing, and imputing genome-wide genotypes in full-sib families when individuals are genotyped with single nucleotide polymorphism (SNP) array or genotyping-by-sequencing (GBS) data. Many applications in genetics and breeding rely on the availability of low-cost high-accuracy genotypes. GBS is an alternative to SNP arrays (Baird et al., 2008; Davey et al., 2011; Elshire et al., 2011), where specific restriction enzymes are used to focus sequencing resources on a limited number of cut sites. GBS is particularly attractive for species without an existing SNP array or as a low-cost alternative to SNP arrays (e.g., Gorjanc et al., 2015, 2017).

GBS data, and in particular low-coverage GBS data, suffers from a large proportion of missing or, when naively called, incorrect genotypes. Unlike SNP array data, where genotypes are called directly from the genotyping platform, with GBS data genotypes must be called from observed sequence reads. It is challenging to accurately call an individual’s genotype when no reads or a small number of reads are generated at a particular locus. Genotype calling accuracy can be increased by considering the haplotypes of other individuals in the population and detecting shared haplotype segments between individuals (Davies et al., 2016; Gorjanc et al., 2017).

Some existing software packages can be used for genotype calling and imputation from GBS data, for example, *Beagle* (Browning and Browning, 2009), *STITCH* (Davies et al., 2016), *AlphaPeel* (Whalen et al., 2018) or *magicimpute* (Zheng et al., 2018). However, these software packages are not designed to exploit the pattern of haplotype sharing observed in large full-sib families. As with traditional imputation methods (e.g., Antolín et al., 2017; O’Connell et al., 2014), we expect that the accuracy of genotype calling, phasing, and imputation from GBS data is highest when population structure is taken into account. In the context of an outbred full-sib family, imputation can be simplified by recognizing that we only need to consider the four parental haplotypes and identify of which pair of haplotypes the offspring inherited at each locus. Here we describe our software package AlphaFamImpute that leverages this particular population structure to improve the accuracy of calling, phasing and imputing genome-wide genotypes and which decreases run-time compared to existing methods. We focus on outbred full-sib families because this represents a population structure commonly found in research populations, and in animal and plant breeding programs.

## Method

AlphaFamImpute performs imputation using a two-step approach. In the first step we call, phase and impute parental genotypes based on the segregation states of their offspring. Segregation states indicate which pair of parental haplotypes an individual inherits at each locus (Ferdosi et al., 2014). We carry out this step iteratively. At each locus, we use the segregation states to project the offspring data to the corresponding parental haplotypes. We combine these parental haplotype estimates with the parents’ data to call parental genotypes at the locus. We then update the offspring segregation states based on the called parental genotypes. Unlike *magicimpute* (Zheng et al., 2018) or *hsphase (Ferdosi et al., 2014)*, we treat the offspring genotype and segregation states probabilistically to account for uncertainty in the genetic data and the called parental haplotypes. In the second step we call, phase, and impute the offspring genotypes by detecting which haplotype segments the offspring inherit from their parents. This process is carried out using multi-locus iterative peeling (Whalen et al., 2018). For a detailed description of the approach, see the Supplementary Information.

Our two-step approach builds closely on previous research. It can be interpreted as: (i) a sampling scheme for the multi-locus iterative peeling (Meuwissen and Goddard, 2010; Whalen et al., 2018); (ii) a probabilistic extension of *hsphase* for full-sib GBS data (Ferdosi et al., 2014); or (iii) an adaptation of *magicimpute* to specifically handle low-coverage GBS data with outbred full-sib individuals (Zheng et al., 2018).

## Software

AlphaFamImpute is written in Python 3 using the *numpy* (Walt et al., 2011) and *numba* (Lam et al., 2015) libraries. It runs on Windows, Linux, and Mac. As inputs, AlphaFamImpute takes in: (i) a genotype file or a sequence read count file, which respectively give the ordered genotypes or sequence read counts for each individual; (ii) a pedigree file which splits the population into full-sib families; and (iii) an optional map file which allows AlphaFamImpute to be run on multiple chromosomes simultaneously. AlphaFamImpute outputs either called genotypes or genotype dosages.

## Example

We demonstrate the performance of AlphaFamImpute on a series of simulated datasets. Each dataset consisted of 100 full-sib families with outbred parents and either 4, 8, 20, 30, 50, or 100 offspring per family. We generated parental haplotypes for 200 parents on a single 100 cM chromosome with 1,000 loci using MaCS (Chen et al., 2009) with an ancestral genetic history set to mimic cattle (Villa-Angulo et al., 2009). We then dropped the haplotypes through the pedigree of full-sib families using AlphaSimR (Gaynor et al., 2019). We generated GBS data by assuming the number of reads at each locus of an individual followed a Poisson distribution with mean equal to a coverage level of 0.5x, 1x, 2x, and 5x and that there was an 0.1% sequencing error rate. The parents either had no GBS data, had low-coverage GBS data at the same coverage as offspring, or had high-coverage (25x) GBS data. We measured imputation accuracy as the correlation between an individual’s true genotype and their imputed genotype dosage averaged across 10 replicates of 100 full-sib families.

Figure 1 presents imputation accuracy across all of the simulations. Imputation accuracy increased with higher GBS coverage, a larger number of genotyped offspring, and more information on the parents. Imputation accuracy was high in a range of cases: if the parents were sequenced at high-coverage imputation accuracy was 0.995 with 15 offspring sequenced at 1x; if the parents were sequenced at the same coverage as the offspring, imputation accuracy was 0.990 with 10 offspring sequenced at 2x; and if the parents had no data, imputation accuracy was 0.997 with 20 offspring sequenced at 2x.

**Figure 1.**
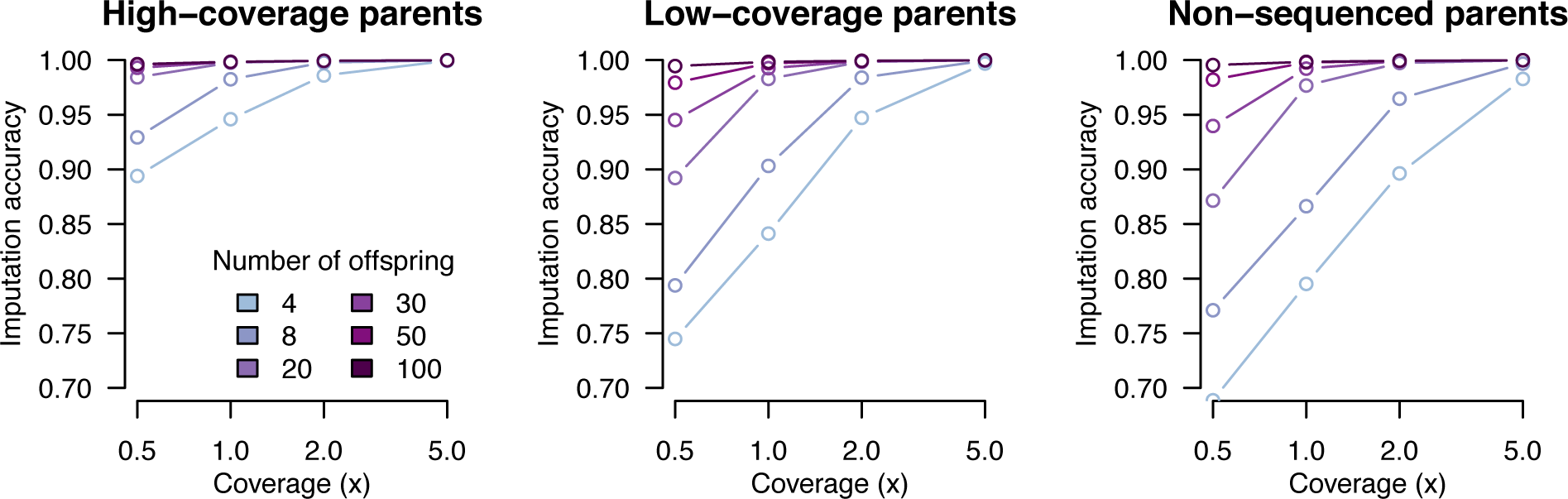
Imputation accuracy for full-sib offspring as a function of sequencing coverage, number of offspring, and parent sequencing coverage.

The primary factor determining imputation accuracy was the total sequencing resources spent on a family. Low sequencing coverage on the parents could be compensated by sequencing additional offspring or sequencing those offspring at higher coverage. When only a few offspring were available this could be compensated by sequencing those offspring at higher coverage.

The computational requirements of AlphaFamImpute were low. When imputing 100 full-sib families with 100 offspring each (total 200 parents and 10,000 offspring) imputation took 106 seconds and used 308 megabytes of memory for 1,000 loci on one chromosome.

## Conclusion

In this paper, we have described the AlphaFamImpute software package for performing fast, high-accuracy calling, phasing and imputing genome-wide genotypes in full-sib families from GBS data. This program will improve the quality of genome-wide genotypes from low-coverage GBS in a range of research and breeding applications.

## Funding and Acknowledgements

The authors acknowledge the financial support from the BBSRC ISPG to The Roslin Institute BB/J004235/1, from Genus PLC, and from Grant Nos. BB/M009254/1, BB/L020726/1, BB/N004736/1, BB/N004728/1, BB/L020467/1, BB/N006178/1 and Medical Research Council (MRC) Grant No. MR/M000370/1.

This work has made use of the resources provided by the Edinburgh Compute and Data Facility (ECDF) (http://www.ecdf.ed.ac.uk).

