## Supplemental Methods for "AlphaFamImpute: high accuracy imputation in full-sib families from genotype-by-sequencing data"

### Supplementary Information for "AlphaFamImpute: high accuracy imputation in full-sib families from genotype-by-sequencing data"

#### Method description

AlphaFamImpute calls, phases and imputes genome-wide genotypes in full-sib families from SNP array and Genotyping-By-Sequencing (GBS) data with a two-stage approach. In the first step it phases and imputes parental genotypes based on the segregation states of their offspring – states which represent which pair of parental haplotypes the offspring inherited. In the second step it calls, phases, and imputes the offspring genotypes by detecting which haplotypes segments the offspring inherit from their parents. In the following we describe generating genotype probabilities from read-count data, the segregation transmission model and the AlphaFamImpute algorithm for calling, phasing and imputing parent and offspring.

##### Genotype probabilities from read counts

In this algorithm we consider diploid individuals genotyped at bi-allelic SNP sites. This results in four possible phased genotype states, aa, aA, Aa, and AA, where 'a' represents a copy of the reference allele, 'A' represents a copy of the alternative allele, the first allele is inherited from the father, and the second allele is inherited from the mother.

With GBS data we assume the input data is sequence read counts for the reference and alternative alleles at each locus of an individual. We follow past work (Xie et al., 2010; Whalen et al., 2018) to translate the observed sequence read counts into genotype probabilities. The probability that an individual, $x$, has genotype $g_{x,i}$ at locus $i$ conditional on observed genetic data $d_{x,i}$ is given by:

$$p(g_{x,i}|d_{x,i}) \propto \begin{matrix} (1-e)^{n_{ref}} e^{n_{alt}} & if\ g_{x,i} = aa \\ \frac{.5^{n_{ref}+n_{alt}}}{2} & if\ g_{x,i} = aA\ or\ Aa \\ (1-e)^{n_{alt}} e^{n_{ref}} & if\ g_{x,i} = AA, \end{matrix} \quad (1)$$

where $n_{ref}$ is the number of sequence reads observed for the reference allele, $n_{alt}$ is the number of sequence reads observed for the alternative allele, and $e$ is a small sequencing error rate,

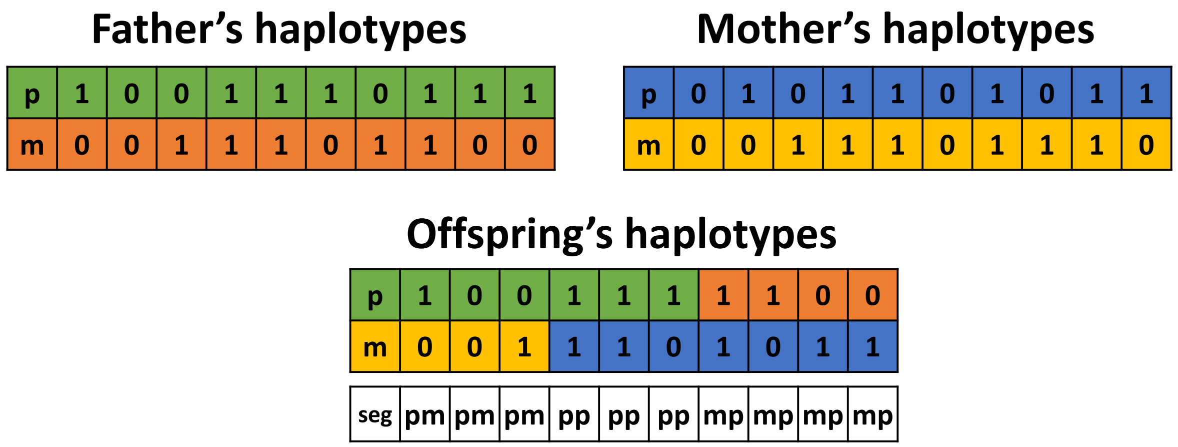

Figure S1: Phased genotypes for a parent-offspring trio. The colors indicate parent haplotypes. In this figure the offspring inherits both grandpaternal (p) and grandmaternal (m) haplotypes from both parents. The "seg" row gives the corresponding segregation state for both of the offspring haplotypes.

assumed to be 0.1% (although it can be changed as a run-time parameter). We also adopt the analogous equations from Whalen et al. (2018) for individuals genotyped with SNP array data.

**Transmission probabilities**

As part of the imputation and phasing algorithms, we track the segregation states for each child. These states represent which pair of parental haplotypes the individual inherits at each locus. We consider four possible segregation states: *pp*, *pm*, *mp*, and *mm*, where the first letter indicates the haplotype the offspring inherited from their father (either the father's paternal (*p*) or father's maternal (*m*) haplotype), and the second letter indicates the haplotype the offspring inherited from their mother. Figure S1 shows haplotypes and segregation states for a parent-offspring trio across 10 loci.

We assume that the segregation states follow a Markov process, with independent recombinations in the maternal and paternal chromosomes. The probability of the segregation state of an offspring, $o$, at locus $i$ conditional on the segregation state at the previous locus, $i+1$ is given by:

$$p(seg_{o,i} = s | seg_{o,i+1} = s') = (1-\gamma)^{2-dif(s,s')}\gamma^{dif(s,s')}, \quad (2)$$

where $dif(s,s')$ is the number of recombinations required to move between state $s$ and state $s'$, for example, if a single recombination occurs between s and s' then:

$$p(seg_{o,i} = pp | seg_{o,i+1} = pm) = (1-\gamma)\gamma.$$

#### Calling, phasing, calling, and imputing the parental genotypes

We call, phase and impute the genotypes of the parents using their genotype probabilities and the genotype and segregation probabilities of their offspring. We start from the last locus on the chromosome and proceed locus-by-locus until the first locus on the chromosome. At each locus we call the genotypes of the parents and re-estimate the segregation probabilities of the offspring based on the called parental genotypes. We describe these steps below.

##### Calling the parental genotypes

We estimate the parental genotypes as the combination of the observed genetic information on the parents with observed genetic information of the offspring. To do this, we first calculate a "posterior" probability term for each offspring (Elston and Stewart, 1971; Kerr and Kinghorn, 1996) which gives the joint parental genotype probabilities of the parents $(g_{f,i}, g_{m,i})$ conditional on the genotype value and the segregation state of an offspring, $o$:

$$posterior_{o,i}(g_{f,i}, g_{m,i}) = \sum_{g_{o,i}} \sum_{seg_{o,i}} p(g_{o,i} | g_{f,i}, g_{m,i}, seg_{o,i}) p(g_{o,i} | d_{o,i}) p(seg_{o,i} | seg_{o,i+1}) \quad (3)$$

The value, $p(g_{o,i} | g_{f,i}, g_{m,i}, seg_{o,i})$, is the probability that a offspring has genotype $g_{o,i}$ conditional on their segregation state, and the genotypes of their parents. This value will either be 0 or 1. As an example, if the segregation state of the offspring is *mm*, where the offspring inherits both of their parents' maternal alleles then:

$$p(g_{o,i} = aa | g_{f,i} = Aa, g_{m.,i} = aa, seg_{o,i} = mm) = 1.$$

Due to possible uncertainty about the underlying genotype state of the offspring and their segregation state, we marginalize over the genotype probabilities for the offspring, $p(g_{o,i}|d_{o,i})$ (calculated from Equation 1), and segregation probabilities, $p(seg_{o,i}|seg_{o,i+1})$ (calculated from Equation 2).

Once the posterior terms are calculated, we evaluate the joint parental genotype probabilities by combining the parent's own genetic information, $p(g_{f,i}|d_{f,i})$ and $p(g_{m,i}|d_{m,i})$, with the genetic information from each offspring:

$$p(g_{f,i}, g_{m,i}) = p(g_{f,i}|d_{f,i})p(g_{m,i}|d_{m,i})\prod_{o} posterior_{o,i}(g_{f,i}, g_{m,i})\,. \tag{4}$$

We called genotypes, $g^*_{m,i}$ and $g^*_{f,i}$, using:

$$g^*_{m,i}, g^*_{f,i} = argmax_{g_{m,i}, g_{f,i}} p(g_{f,i}, g_{m,i}). \tag{5}$$

If there are multiple genotype states that satisfy Equation 5, then we select one of those states uniformly at random.

**Updating the offspring segregation probabilities**

Once we call parental genotypes at a particular locus, we update the offspring segregation probabilities. We do this update by combining a right-hand segregation probability estimate $p(seg_{o,i}|seg_{o,i+1})$, with the conditional probability of an individual's segregation state conditional on the parental genotypes, $p(seg_{o,i}|g^*_{m,i}, g^*_{f,i})$:

$$p(seg_{o,i}|g^*_{m,i}, g^*_{f,i}, seg_{o,i+1}) \propto p(seg_{o,i}|g^*_{m,i}, g^*_{f,i})\; p(seg_{o,i}|seg_{o,i+1}) \tag{6}$$

The probability of an individual's segregation state conditioned on the genotype state of their parents is:

$$p\left(seg_{o,i} \middle| g_{m,i}^*, g_{f,i}^*\right) \propto \sum_{g_{o,i}} p\left(g_{o,i} \middle| g_{m,i}^*, g_{f,i}^*, seg_{o,i}\right) p\left(g_{o,i} \middle| d_{o,i}\right), \tag{7}$$

where $p\left(g_{o,i} \middle| g_{m,i}^*, g_{f,i}^*, seg_{o,i}\right)$ is as in Equation 3, and $p\left(g_{o,i} \middle| d_{o,i}\right)$ is given by Equation 1.

We calculate the right-hand estimate, $p\left(seg_{o,i} \middle| seg_{o,i+1}\right)$ by marginalizing over segregation states at locus $i + 1$:

$$p\left(seg_{o,i} \middle| seg_{o,i+1}\right) = \sum_{s'} p\left(seg_{o,i} = s \middle| seg_{o,i+1} = s'\right) p\left(seg_{o,i+1} \middle| g_{m,i}^*, g_{f,i}^*, seg_{o,i+2}\right), \tag{8}$$

where $p\left(seg_{o,i} = s \middle| seg_{o,i+1} = s'\right)$ is given by Equation 2. Equation 8 requires the recursive calculation of $p\left(seg_{o,i+1} \middle| g_{m,i}^*, g_{f,i}^*, seg_{o,i+2}\right)$, which can be calculated in linear time by storing the values at each locus.

**Additional details**

We made two minor modifications to this approach in order to increase accuracy. First, we phase the parental genotypes twice: in a backward pass where the initial segregation probabilities are set to a uniform value, $p\left(seg_{o,N}\right) = .25$ (where N is the number of loci), and in a forward pass where the segregation probabilities at the first locus are based on the final segregation probabilities from the backward pass:

$$p^*\left(seg_{o,1}\right) = p\left(seg_{o,1} \middle| seg_{o,2}, g_{m,1}^*, g_{f,1}^*\right) \tag{8}$$

where the asterisk (*) denotes that this is the probability for the forward pass. Working in a two-pass approach allows us to correctly handle uncertainty in the segregation states at the start of the backward pass, while still maintaining high accuracy at both ends of the chromosome. Second, we avoid underflow in Equation 1 and Equation 4 by calculating and combining the genotype probability estimates on a log-scale, as done by Kerr and Kinghorn (1996), for example.

#### Phasing and imputing the offspring

We use multi-locus peeling to call, impute, and phase the offspring. This method is equivalent to using a diploid hidden Markov model where the parental haplotypes form the reference libraries for each of the offspring's haplotypes (Li and Stephens, 2003). We calculate the probability of an individual's genotype based on their own genetic data, and the called parental genotypes conditional on the individual's segregation probabilities:

$$p(g_{o,i}) = p(g_{o,i}|d_{o,i}) \sum_{seg_{o,i}} p(g_{o,i}|g^*_{m,i}, g^*_{f,i}, seg_{o,i}) p(seg_{o,i}|seg_{o,i+1}) p(seg_{o,i}|seg_{o,i-1}), \quad (9)$$

where $p(g_{o,i}|d_{o,i})$ is given by Equation 1 and we calculate the left-hand and right-hand segregation probabilities, $p(seg_{o,i}|seg_{o,i+1})$ and $p(seg_{o,i}|seg_{o,i-1})$, recursively using Equation 8.

Based on past experience, we have found that incorrectly called parental genotypes can reduce imputation accuracy. In order to mitigate this issue, we only evaluate the segregation probabilities in Equation 8 based on loci where $p(g^*_{f,i}, g^*_{m,i}) > .95$. We also modify Equation 9 to marginalize across parental genotypes states, replacing $p(g_{c,i}|g^*_{m,i}, g^*_{f,i}, seg_{c,i})$ with

$$\sum_{g_{m,i}, g_{f,i}} p(g_{c,i}|g_{m,i}, g_{f,i}, seg_{c,i}) p(g_{m,i}, g_{f,i}). \quad (10)$$
